# Addressing Gaps in HIV Preexposure Prophylaxis Care to Reduce Racial Disparities in HIV Incidence in the United States

**DOI:** 10.1101/249540

**Authors:** Samuel M. Jenness, Kevin M. Maloney, Dawn K. Smith, Karen W. Hoover, Steven M. Goodreau, Eli S. Rosenberg, Kevin M. Weiss, Albert Y. Liu, Darcy W. Rao, Patrick S. Sullivan

**Author notes:** Corresponding Author: Emory University, 1520 Clifton Road, Atlanta, GA 30322.

## Abstract

The potential for HIV preexposure prophylaxis (PrEP) to reduce the racial disparities in HIV incidence in the United States may be limited by racial gaps in PrEP care. We used a network-based mathematical model of HIV transmission for younger black and white men who have sex with men (B/WMSM) in the Atlanta area to evaluate how race-stratified transitions through the PrEP care continuum from initiation to adherence and retention could impact HIV incidence overall and disparities in incidence between races, using current empirical estimates of BMSM continuum parameters. Relative to a no-PrEP scenario, implementing PrEP according to observed BMSM parameters was projected to yield a 23% decline in HIV incidence (HR = 0.77) among BMSM at year 10. The racial disparity in incidence in this observed scenario was 4.95 per 100 person-years at risk (PYAR), a 19% decline from the 6.08 per 100 PYAR disparity in the no-PrEP scenario. If BMSM parameters were increased to WMSM values, incidence would decline by 47% (HR = 0.53), with an associated disparity of 3.30 per 100 PYAR (a 46% decline in the disparity). PrEP could simultaneously lower HIV incidence overall and reduce racial disparities despite current gaps in PrEP care. Interventions addressing these gaps will be needed to substantially decrease disparities.

---

HIV prevalence among black men who have sex with men (BMSM) is 3–6 times as high as white MSM (WMSM) across the United States, with incidence increasing among younger BMSM (1,2). The causes of these disparities have been challenging to quantify. Although HIV medical care engagement has been worse for BMSM (3), behavioral studies consistently suggest lower HIV acquisition risks for BMSM than WMSM (4,5). The US National HIV/AIDS Strategy has among its goals to reduce both new HIV diagnoses by 25% overall and racial disparities in diagnoses by 15% by 2020 (6), with several strategies prioritized.

One high-priority intervention is scaling-up HIV preexposure prophylaxis (PrEP), which has proven highly effective at lowering HIV risk (7). Yet it is uncertain whether PrEP can be used to reduce HIV racial disparities. PrEP use by MSM has increased nationally since FDA approval. Pharmacy data indicate a 500% increase in PrEP prescriptions since 2014, but black persons received only 10% of those despite accounting for nearly half of recent HIV diagnoses (2,8). Open-label PrEP studies have consistently highlighted challenges in reaching BMSM (9–19). Reducing racial disparities in HIV incidence could be achieved with PrEP as part of a comprehensive HIV prevention approach (20), but whether that is possible given the major gaps in PrEP care for BMSM remains a critical unanswered question.

A PrEP care continuum framework conceptually defines these gaps. Kelley et al., for example, identified the steps towards complete HIV prevention with PrEP via awareness of PrEP, access to PrEP-related healthcare services, obtaining a PrEP prescription, and adherence after initiation (21). Their race-stratified estimates, based on data from an HIV cohort in Atlanta (22), suggested that BMSM had equal or worse outcomes on all four steps. Nunn et al. included a fifth step: retention in PrEP care after effective adherence (23). Although a continuum framework does not directly solve the problem of *how* to close these gaps, it organizes research priorities and prevention efforts into distinct targets for intervention.

In this study, we used mathematical modeling to 1) quantify the PrEP-related reduction in HIV incidence for younger BMSM in the Southeastern US over the next decade given current PrEP care continuum estimates; and 2) predict how improvements along each continuum step (awareness, access, prescription, adherence, and retention) for BMSM, individually and jointly, could further reduce HIV incidence overall and disparities in HIV incidence between races. Although the levels of HIV disparities and scale-up of PrEP vary across health jurisdictions and risk groups in the US, findings from this high-burden, low-resource target population may broadly inform intervention strategies through which PrEP could meet current HIV disparity reduction goals nationally.

## METHODS

We previously developed a mathematical model for HIV/STI transmission dynamics for US MSM using the *EpiModel* software platform (24), a computational toolkit for simulating epidemics over dynamic sexual networks under the statistical framework of temporal exponential random graph models (TERGMs) (25). Our prior applications investigated the sources of HIV racial disparities among MSM in Atlanta and the potential impact of PrEP for MSM across races (26,27). This study integrated these two research streams to develop the model structure, parameterization, and analyses for simulating PrEP stratified by race and represent PrEP care on a continuum framework. Full methodological details are provided in an Appendix.

### HIV Transmission and Progression

Our model simulates the dynamics of main, casual, and one-time sexual partnerships for non-Hispanic BMSM and WMSM, aged 18–40 (26,27). Predictors of partnership formation included partnership type, number of ongoing partnerships, race and age mixing, and sorting by receptive versus insertive sexual position. For main and casual partnerships, we modeled relational dissolution as a constant hazard reflecting their median durations. All network model terms were stratified by race.

MSM progressed through HIV disease in the absence of antiretroviral therapy (ART) with evolving HIV viral loads that modified the rate of HIV transmission (28). After infection, men were assigned into clinical care trajectories controlling rates of HIV diagnosis, ART initiation, and HIV viral suppression (29,30). ART was associated with decreased mortality and lower HIV transmissibility (31). Other factors modifying the HIV acquisition probability included current infection with other sexually transmitted infections (32), condom use (33), sexual position (34), and circumcision of the insertive partner (35).

Parameters for network/behavioral features of the model were estimated from two studies of HIV disparities between younger BMSM and WMSM in Atlanta, our target population (22,36). Involvement was a prospective HIV incidence cohort (n=803) and the MAN Project was a cross-sectional chain-referral sexual network study (n = 314). Venue-time-space sampling was used for both studies to minimize selection biases. Remaining model parameters for the underlying model (e.g., HIV natural history and ART clinical effects) were assumed to be common across MSM populations and therefore drawn from secondary literature sources. Methods for data analysis and assumptions for model parameterization are described in greater detail in the Appendix and Goodreau et al. (26).

### PrEP Continuum

We represented PrEP based on a five-step continuum: awareness of PrEP, access to healthcare, likelihood of receiving a prescription, effective adherence, and retention in care. Race-stratified probabilities governing transitions across steps were drawn from two PrEP demonstration projects (11,21). Awareness was estimated as 50% for both races, whereas access was 76% for BMSM and 95% for WMSM. Both were fixed attributes assigned at entry into the network.

Prescription probabilities were 63% for BMSM and 73% WMSM, simulated as a Bernoulli random draw at the point of clinical evaluation for and precondition of initiating PrEP: diagnostic HIV screening. Screening rates were stratified by race based on empirical data (see Appendix), but were assumed homogenous otherwise. Consistent with prior models (32), we simulated the four biobehavioral indications for starting PrEP defined in the CDC guidelines (37): higher-risk sexual behavior in various partnership configurations or an STI diagnosis within the prior 6 months. Because indications were time-varying, the probability of a PrEP prescription was therefore a joint function of the race-specific probability of receiving a prescription plus current indications at HIV screening.

Effective PrEP adherence in the model represented men taking 4+ doses per week across follow-up (11). Proportions meeting this criterion were 60% for BMSM and 93% for WMSM. Taking PrEP at this dosage was been associated with a 98% relative reduction in HIV acquisition risk per sexual act, following Grant et al (38). MSM who were adherent to PrEP at this level reduced their condom use with by 40% (39). PrEP discontinuation (the converse of retention) rates were based on observed proportions of MSM with indications who had stopped PrEP by week 48 of follow-up (43.8% for BMSM and 18.3% for WMSM). We transformed these proportions into median times to discontinuation (1.1 years and 3.2 years, respectively) assuming a hypergeometric distribution. We simulated this form of spontaneous discontinuation conditional on having ongoing indications, consistent with our data analysis. In addition, men stopped PrEP if they no longer exhibited PrEP indications (evaluated annually for active PrEP users) (37).

### Counterfactual Scenarios

To estimate the causal impact of changes to the PrEP continuum for BMSM, we varied the probabilities for each of the five steps individually and jointly. The reference scenario to which all intervention counterfactuals were compared was that no MSM (of either race) were on PrEP. While this scenario does not represent a proposed public health strategy, it provides maximum analytical clarity for estimating HIV disparities before and after the introduction of PrEP. Furthermore, we calibrated the model to race-stratified HIV prevalence estimates in 2013, just after the FDA approval of PrEP (37).

For individual continuum steps, we set the parameters for BMSM to those marginal values observed for WMSM and then higher levels, while holding other BMSM continuum parameters fixed at their observed levels. The WMSM continuum and all other model parameters, including those governing risk behavior and HIV clinical care, were always held fixed across all scenarios; results here are conditional on that assumption. For scenarios modifying parameters in combination, we varied the BMSM parameters on a relative scale. Scenarios in which BMSM parameters were set to 150% of observed values, for example, multiplied each of the empirical estimates by 1.5 (with individual probabilities capped at 1). For our final analysis, we grouped the five continuum steps into two factor groups — initiation (awareness, access, and prescription) and engagement (adherence and retention) — and then projected outcomes across a spectrum of relative BMSM values in each group.

### Calibration, Simulation, and Analysis

With a starting network size of 10,000 MSM (aged 18–40), 50% were initialized in each race, a ratio that approximates the distribution for the Atlanta area and provides analytical clarity (26). We calibrated our model to observed race-specific HIV prevalence at baseline in an Atlanta-based cohort: 43.4% for BMSM and 13.2% for WMSM (22). Based on prior work on modeling the causes of these disparities (26), we incorporated the full 95% confidence intervals of estimated rates of anal intercourse and probabilities of condom use for model calibration. We also implemented race-specific parameters simulating condom failure (due to slippage or breakage), consistently higher in BMSM (40–42), and diagnostic screening for bacterial STIs (increasing the risk of HIV if untreated), often lower for BMSM (4). Approximate Bayesian computation methods estimated the values of these parameters best fitting the observed prevalence data (32,43). The calibrated model provided an excellent fit to these targets. We also successfully externally validated this calibration with an “out-of-model” prediction of HIV prevalence by the interaction of race and age (see Appendix Section 12).

Intervention models simulated each scenario over a 10-year time horizon. For each scenario, we simulated the model 250 times and summarized the distribution of results based on median values and 95% credible intervals (CrI). Outcomes were race-specific HIV prevalence and incidence per 100 person years at risk (PYAR), and the hazard ratio comparing incidence to the no-PrEP reference scenario, all at year 10. The percent of infections averted (PIA) among BMSM compared the cumulative incidence in each intervention scenario to that of the reference scenario. The number needed to treat (NNT) was the number of BMSM person-years on PrEP required to avert one new HIV infection for BMSM. Two disparity indices were calculated to compare PrEP impact for BMSM versus WMSM: the absolute disparity was the difference in incidence rates for BMSM and WMSM, and the relative disparity was the ratio of those rates. Finally, we calculated a prevention index as the difference in hazard ratios associated with PrEP uptake for BMSM and WMSM.

## RESULTS

Table 1 shows the impact of individual PrEP continuum steps for BMSM at observed and counterfactual values. In comparison with the reference scenario in which no one (of either race) received PrEP, the observed BMSM PrEP continuum scenario projected 8.4% (CrI = 7.7, 9.1) of BMSM to be on PrEP across follow-up. This yielded a 3-percentage point decline in HIV prevalence (39.9% versus 43.1%) and a 23% decline in incidence (HR = 0.77; CrI = 0.57, 0.99) among BMSM at year 10. The cumulative PIA was 14.1% (CrI = 8.2%, 21.0%) for BMSM over the intervention horizon. We then modeled changes to individual steps (holding parameters for remaining steps at observed BMSM values). For awareness, while the observed values were equal for both races, increasing that awareness proportion for BMSM had a strong impact on PrEP use, and with that, declines in incidence. For access, setting the BMSM access parameter to the observed WMSM value (95%) resulted in a smaller decline in incidence than changes to awareness. Conditional on access, empirical differences in the probability of prescription were relatively small. Increasing the proportion highly adherent did not impact the overall proportion of BMSM on PrEP (which includes PrEP users across adherence levels); this also resulted in a relatively small prevention effect. Higher levels of retention on PrEP were associated with greater PrEP prevention benefits because fewer MSM indicated for PrEP were cycled off PrEP during their periods of high sexual risk.

**Table 1.** Proportion on HIV Preexposure Prophylaxis (PrEP), HIV Prevalence and Incidence per 100 PYAR at Year 10, Hazard Ratio of Incidence at Year 10 and Percent of Infections Averted (PIA) over 10 Years Compared to the Reference (No PrEP) Scenario among Black MSM, by PrEP Continuum Step Values

| Scenario | % BMSM<br>On PrEP<br>95% CrI<sup>3</sup> | BMSM HIV<br>Prevalence<br>95% CrI | BMSM HIV<br>Incidence<br>95% CrI | BMSM Hazard<br>Ratio<br>95% CrI | BMSM PIA<br>95% CrI |
| --- | --- | --- | --- | --- | --- |
| **Reference (No PrEP)** | 0.0 (0.0, 0.0) | 43.1 (41.2, 46.3) | 7.73 (6.51, 9.07) | 1.00 | - |
| **Aware of PrEP (%)** |  |  |  |  |  |
| 30% | 5.0 (4.3, 5.5) | 41.8 (39.8, 43.8) | 6.54 (5.47, 7.68) | 0.85, 0.68, 1.09) | 9.3 (1.8, 15.3) |
| 50% (Obs B & W)<sup>1,2</sup> | 8.4 (7.7, 9.1) | 39.9 (38.1, 41.6) | 5.88 (4.66, 7.05) | 0.77 (0.57, 0.99) | 14.1 (8.2, 21.0) |
| 70% | 12.0 (11.2, 12.8) | 37.7 (35.7, 39.5) | 5.24 (4.43, 6.25) | 0.68 (0.55, 0.88) | 20.0 (13.8, 26.9) |
| 90% | 15.6 (14.8, 16.5) | 35.9 (34.2, 37.8) | 4.68 (3.88, 5.57) | 0.61 (0.48, 0.78) | 25.0 (18.7, 30.4) |
| **Access to PrEP Given Awareness** |  |  |  |  |  |
| 50% | 5.4 (4.9, 5.9) | 41.4 (39.6, 43.3) | 6.43 (5.30, 7.52) | 0.84 (0.65, 1.06) | 10.1 (3.3, 16.4) |
| 76% (Obs B)<sup>1</sup> | 8.4 (7.7, 9.1) | 39.9 (39.1, 41.6) | 5.88 (4.66, 7.05) | 0.77 (0.57, 0.99) | 14.1 (8.2, 21.0) |
| 85% | 9.4 (8.8, 10.1) | 39.1 (37.5, 41.0) | 5.65 (4.73, 6.81) | 0.73 (0.58, 0.95) | 15.9 (10.3, 22.4) |
| 95% (Obs W)<sup>2</sup> | 10.6 (9.9, 11.5) | 38.5 (36.3, 40.3) | 5.54 (4.56, 6.48) | 0.72 (0.56, 0.94) | 18.2 (10.8, 24.9) |
| **Prescribed PrEP Given Access** |  |  |  |  |  |
| 50% | 7.2 (6.5, 7.8) | 40.4 (38.4, 42.2) | 6.14 (5.13, 7.02) | 0.81 (0.64, 0.99) | 12.9 (4.8, 18.5) |
| 63% (Obs B)<sup>1</sup> | 8.4 (7.7, 9.1) | 39.9 (38.1, 41.6) | 5.88 (4.66, 7.05) | 0.77 (0.57, 0.99) | 14.1 (8.2, 21.0) |
| 73% (Obs W)<sup>2</sup> | 9.2 (8.5, 9.9) | 39.5 (37.6, 41.3) | 5.83 (4.84, 6.90) | 0.75 (0.59, 0.98) | 15.4 (8.8, 21.7) |
| 85% | 10.1 (9.4, 10.9) | 38.9 (36.8, 40.5) | 5.64 (4.54, 6.73) | 0.74 (0.54, 0.95) | 17.3 (9.5, 23.4) |
| **Effective Adherence Given Prescription** |  |  |  |  |  |
| 50% | 8.3 (7.7, 8.9) | 39.8 (38.1, 41.8) | 5.98 (4.78, 7.04) | 0.78 (0.58, 1.02) | 14.2 (6.0, 20.5) |
| 60% (Obs B)<sup>1</sup> | 8.4 (7.7, 9.1) | 39.9 (38.1, 41.6) | 5.88 (4.66, 7.05) | 0.77 (0.57, 0.99) | 14.1 (8.2, 21.0) |
| 75% | 8.6 (7.8, 9.2) | 39.3 (37.2, 41.1) | 5.70 (4.61, 6.73) | 0.75 (0.56, 0.93) | 15.6 (9.9, 23.3) |
| 93% (Obs W)<sup>2</sup> | 8.8 (8.1, 9.4) | 38.9 (37.0, 40.6) | 5.59 (4.66, 6.78) | 0.74 (0.58, 0.95) | 16.5 (10.1, 22.6) |
| **Time on PrEP before Discontinuation Given Prescription** |  |  |  |  |  |
| 1 year | 7.9 (7.2, 8.6) | 40.1 (37.9, 42.0) | 5.97 (4.89, 7.02) | 0.78 (0.60, 0.99) | 13.7 (7.6, 20.0) |
| 1.1 years (Obs B)<sup>1</sup> | 8.4 (7.7, 9.1) | 39.9 (38.1, 41.6) | 5.88 (4.66, 7.05) | 0.77 (0.57, 0.99) | 14.1 (8.2, 21.0) |
| 2 years | 10.7 (9.9, 11.4) | 38.6 (36.8, 40.6) | 5.45 (4.41, 6.69) | 0.71 (0.54, 0.93) | 17.5 (11.0, 23.7) |
| 3.16 years (Obs W)<sup>2</sup> | 12.3 (11.6, 13.0) | 37.8 (35.6, 39.4) | 5.23 (4.11, 6.22) | 0.68 (0.52, 0.87) | 20.0 (13.5, 26.8) |

<sup>1</sup> Observed PrEP Continuum value for Black MSM, serving as reference value for step when parameters for other steps were modified in counterfactuals. <sup>2</sup> Observed PrEP Continuum value for White MSM. <sup>3</sup> CrI = credible intervals.

In Table 2, we project the impact of scaling the BMSM continuum parameters jointly on HIV incidence outcomes for both BMSM and WMSM. Compared to the scenario in which all BMSM continuum parameters were set to observed levels for BMSM, when all BMSM parameters were set to levels observed for WMSM we project that 17.7% (CrI = 16.8%, 18.7%) of BMSM would actively be on PrEP. This compares to 23.4% (CrI = 22.4%, 24.4%) of WMSM, with the difference due to WMSM’s higher level of CDC PrEP indications even when all continuum parameters were equal. In this scenario where BMSM parameters were set to observed WMSM values, incidence among BMSM would be lower (HR = 0.53 versus 0.77) than in the scenario in which BMSM parameters were set to observed BMSM values. Scaling up BMSM continuum parameters to even higher levels (150% or 200% of observed BMSM values) would result in even greater numbers of BMSM on PrEP, with stronger incidence reductions BMSM. Overall, all levels of PrEP modeled (even those poorer than observed) resulted in a reduction in HIV incidence for BMSM compared to no PrEP, with increasing initiation and engagement associated with incidence declines by greater than three-quarters at optimistic implementation levels.

**Table 2.** Proportion of MSM on HIV Preexposure Prophylaxis (PrEP), HIV Incidence per 100 PYAR at Year 10, Hazard Ratio of Incidence at Year 10 Compared to the Reference (No PrEP) Scenario, by Race, and Disparity Indices, Across Combined Relative PrEP Continuum Indicator Scenarios for Black MSM

| Scenario | Black MSM Outcomes |  |  | White MSM Outcomes |  |  | Disparity Indices |  |  |
| --- | --- | --- | --- | --- | --- | --- | --- | --- | --- |
|  | % On PrEP<br>95% CrI<sup>4</sup> | HIV Incidence<br>95% CrI | Hazard Ratio<br>95% CrI | % On PrEP<br>95% CrI | HIV Incidence<br>95% CrI | Hazard Ratio<br>95% CrI | Absolute<sup>1</sup><br>95% CrI | Relative<sup>2</sup><br>95% CrI | Prevention<sup>3</sup><br>95% CrI |
| **Reference (No PrEP)** | 0.0 (0.0, 0.0) | 7.73 (6.51, 9.07) | 1.00 | 0.0 (0.0, 0.0) | 1.65 (1.27, 2.07) | 1.00 | 6.08 | 4.68 | - |
| **Combined PrEP Continuum for BMSM** |  |  |  |  |  |  |  |  |  |
| Obs. B x 50% Rel. | 0.8 (0.6, 0.9) | 7.38 (6.20, 8.69) | 0.97 (0.74, 1.19) | 23.4 (22.3, 24.5) | 1.00 (0.69, 1.32) | 0.61 (0.40, 0.88) | 6.38 | 7.38 | 0.36 |
| Obs. B | 8.4 (7.7, 9.1) | 5.88 (4.66, 7.05) | 0.77 (0.57, 0.99) | 23.4 (22.4, 24.4) | 0.93 (0.68, 1.23) | 0.56 (0.40, 0.86) | 4.95 | 6.32 | 0.21 |
| Obs. W | 17.7 (16.8, 18.7) | 4.15 (3.36, 4.98) | 0.53 (0.43, 0.70) | 23.4 (22.2, 24.5) | 0.85 (0.59, 1.15) | 0.52 (0.34, 0.79) | 3.30 | 4.88 | 0.01 |
| Obs. B x 150% Rel. | 27.2 (26.1, 28.4) | 3.06 (2.33, 3.80) | 0.39 (0.29, 0.51) | 23.5 (22.4, 24.5) | 0.77 (0.47, 1.10) | 0.47 (0.28, 0.73) | 2.29 | 3.97 | -0.08 |
| Obs. B x 200% Rel. | 42.1 (40.8, 43.4) | 1.73 (1.33, 2.17) | 0.22 (0.17, 0.32) | 23.5 (22.4, 24.6) | 0.69 (0.48, 0.99) | 0.42 (0.27, 0.66) | 1.04 | 2.51 | -0.2 |

<sup>1</sup> Absolute disparity index = BMSM HIV incidence - WMSM HIV incidence. <sup>2</sup> Relative disparity index = BMSM HIV incidence / WMSM HIV incidence. <sup>3</sup> Prevention index = BMSM hazard ratio - WMSM hazard ratio. <sup>4</sup> CrI = credible intervals.

Figure 1 graphically depicts this relative scaling of the joint BMSM parameters. Changes in outcomes are non-linear over these relative parameter changes, with the greatest marginal gains from scaling up the parameters in the range of 1.0 to 1.5 of observed. Although we never modified the WMSM PrEP continuum parameters (see Table 2, with the proportion on PrEP stable in all scenarios), WMSM incidence declined from 0.93 (CrI = 0.68, 1.23) per 100 PYAR in the observed BMSM scenario to 0.69 (CrI = 0.48, 0.99) in the 200% scenario. These are all indirect effects from BMSM PrEP use, possible because 11% of sexual partnerships on average were between-race.

**Figure 1.**
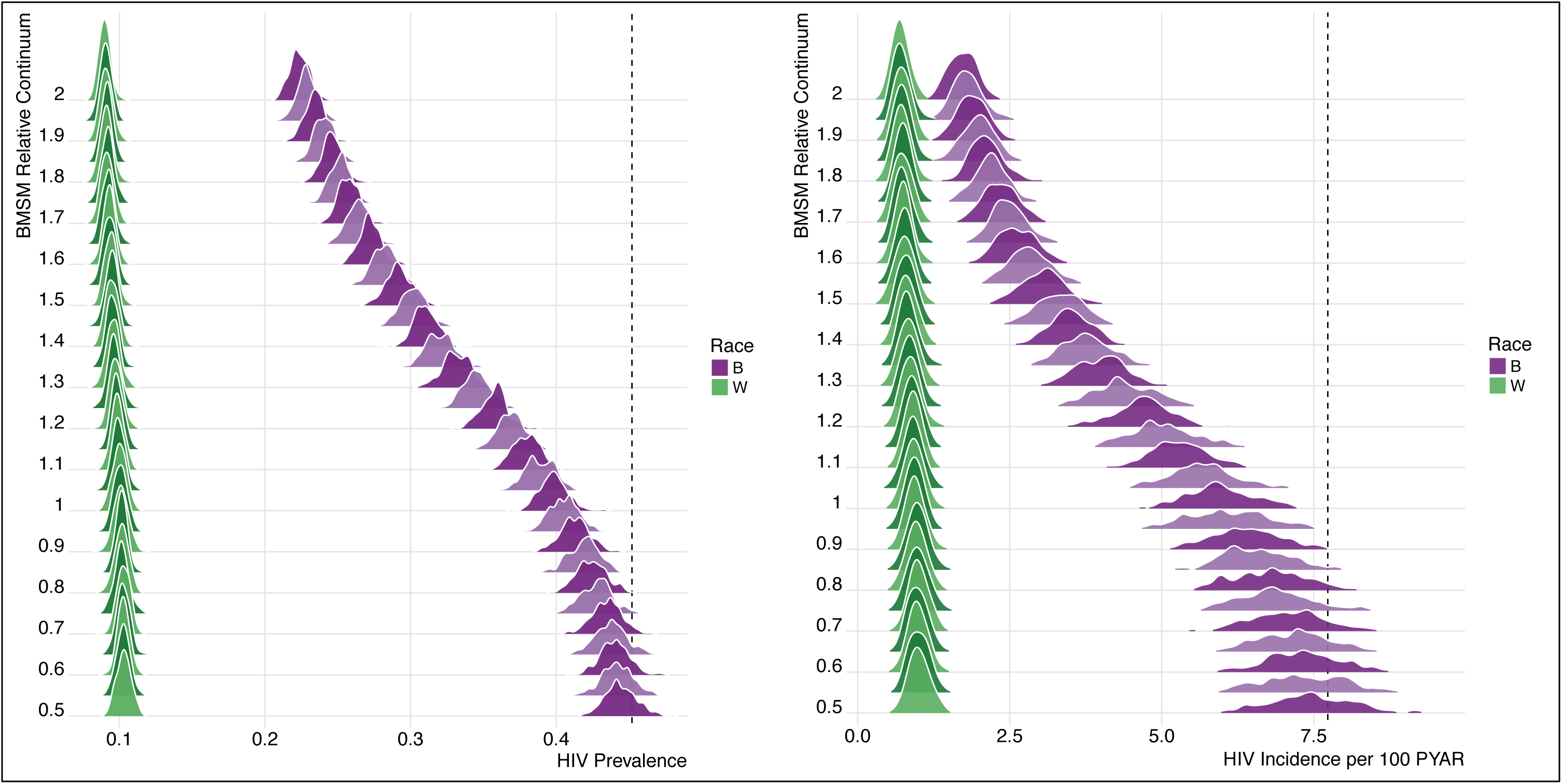
Empirical distribution of model simulations (n = 250 in each scenario) for race-specific HIV prevalence and HIV incidence (per 100 person-years at risk) at Year 10 for BMSM and WMSM across relative values of the combined BMSM PrEP continuum (awareness, access, prescription, adherence, and retention). Relative value 1.0 is the observed BMSM continuum values, 0.5 is half of those observed, and 2.0 is twice those observed. The vertical dashed line indicates HIV prevalence and incidence among BMSM in the reference “no PrEP” scenario.

The impact of PrEP on HIV disparities is also shown in Table 2 and Figure 2. The absolute disparity in the no-PrEP scenario was 6.08 per 100 PYAR, depicted by the dashed horizontal line. Each dot in the figure represents one simulation, across the range of simulated relative BMSM continuum values (0.5– 2.0). The set of points at a given x-axis value therefore represents uncertainty in the relationship between the continuum value and disparity measure as function of the inherent stochastic variation in the model. Implementing PrEP under the observed BMSM scenario (dotted vertical line) would reduce the absolute disparity compared to the scenario with no PrEP (4.95 per 100 PYAR), a 19% decline. If BMSM parameters were set to observed WMSM values, incidence would decline by 47% (HR = 0.53) among BMSM, with an absolute disparity of 3.30 per 100 PYAR, a 46% decline. The prevention index, the difference in hazard ratios, was effectively zero (0.01) in the scenario with BMSM parameters set to WMSM values, and even lower (indicating a greater individual-level prevention effect for BMSM) as the continuum is scaled up. Reductions in the absolute disparity index coincide with reductions in the prevention index, however, parity in the hazard ratios by race (i.e., the same individual-level effect of PrEP) is not necessary to reduce absolute disparities (i.e., population-level difference in incidence).

**Figure 2.**
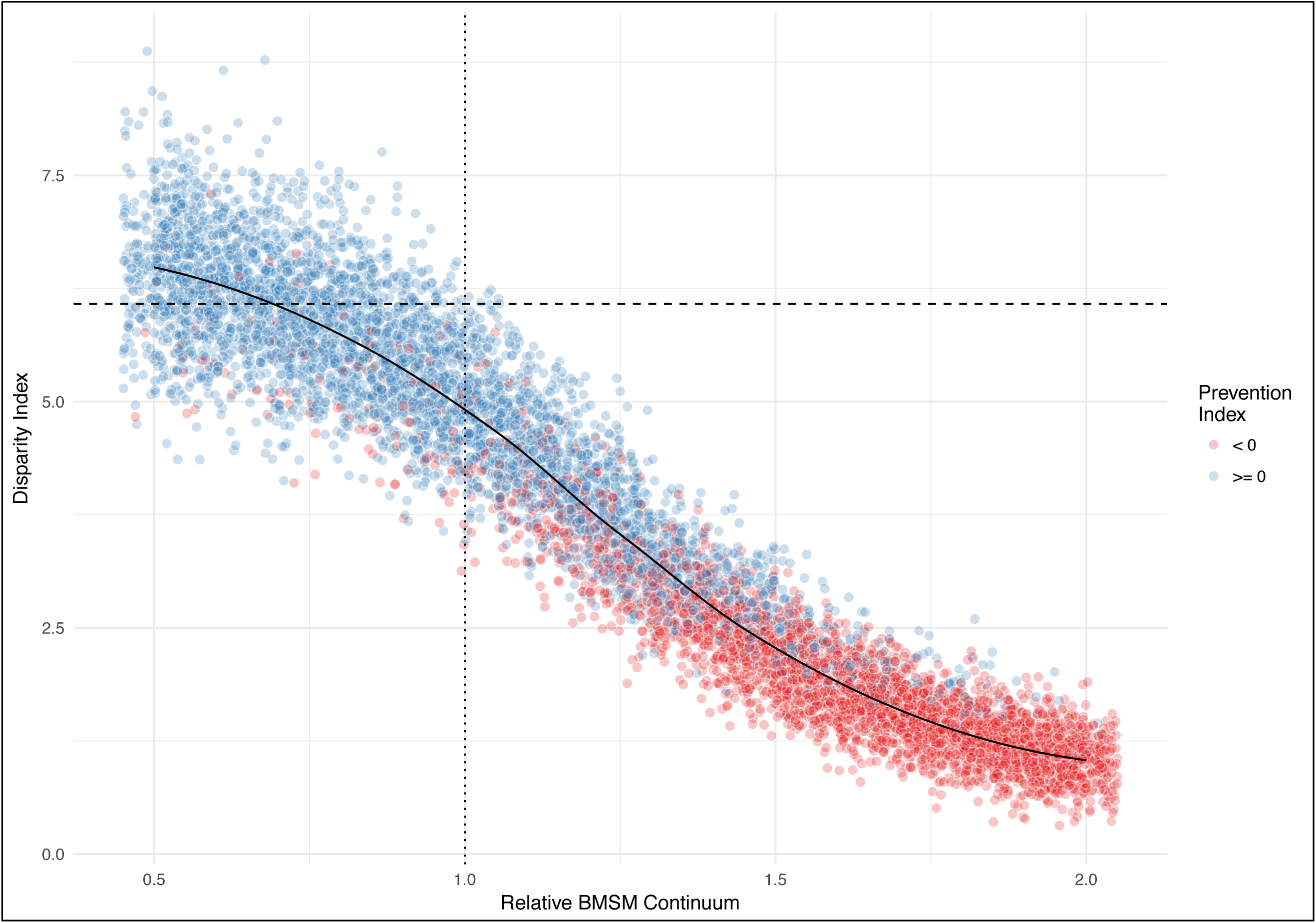
The absolute disparity index (HIV incidence in BMSM - HIV incidence in WMSM) and Prevention Index (HR from HIV preexposure prophylaxis [PrEP] for black MSM [BMSM] - HR from PrEP for white MSM [WMSM]) across relative values of the combined BMSM PrEP continuum, at year 10. Dashed horizontal line shows the pre-PrEP disparity index; dotted vertical line shows the empirical BMSM continuum values. Each dot represents one simulation. Dots were slightly horizontally jittered to reduce over-plotting.

The relative disparity index tells a different story. Under no PrEP, the predicted relative disparity was 4.68, whereas the disparity would increase to 6.32 in the observed BMSM scenario. Relative disparities increased despite higher PrEP use among BMSM because this relative measure is sensitive to changes in its denominator (i.e., WMSM incidence, as a function of their PrEP use). Only when the individual-level benefit of PrEP is greater for BMSM compared to WMSM (i.e., the prevention index is less than 0) do the relative disparities fall below levels in the no-PrEP scenario. Overall levels of effective PrEP care for BMSM would need to be greater or equal to those for WMSM to generate a reduction in the disparity on a relative scale.

Figure 3 aggregates the PrEP continuum into two factor groups of initiation (awareness, access, and prescription) and engagement (adherence and retention), with counterfactual levels of BMSM PrEP parameters in each group and outcomes of BMSM PIA and NNT. In the left panel, greater gains in the PIA for BMSM are projected with an increase in the initiation factors (moving left to right) compared to the same proportional increase in the engagement factors (moving bottom to top), shown by the relatively vertical orientation of the bands at the 1.0/1.0 intersection. At worse than observed initiation levels, little is gained by improving engagement. In the right panel the NNT at observed initiation factor levels ranges from approximately 9 to 13 years of BMSM person-time on PrEP to prevent one new BMSM infection. The NNT is lower as engagement is scaled up because adherence increases the per-dose prevention efficiency. The NNT is higher as initiation factors are scaled up because high PrEP coverage leads to substantial declines in the HIV incidence rate, requiring more person-time on PrEP to prevent an infection.

**Figure 3.**
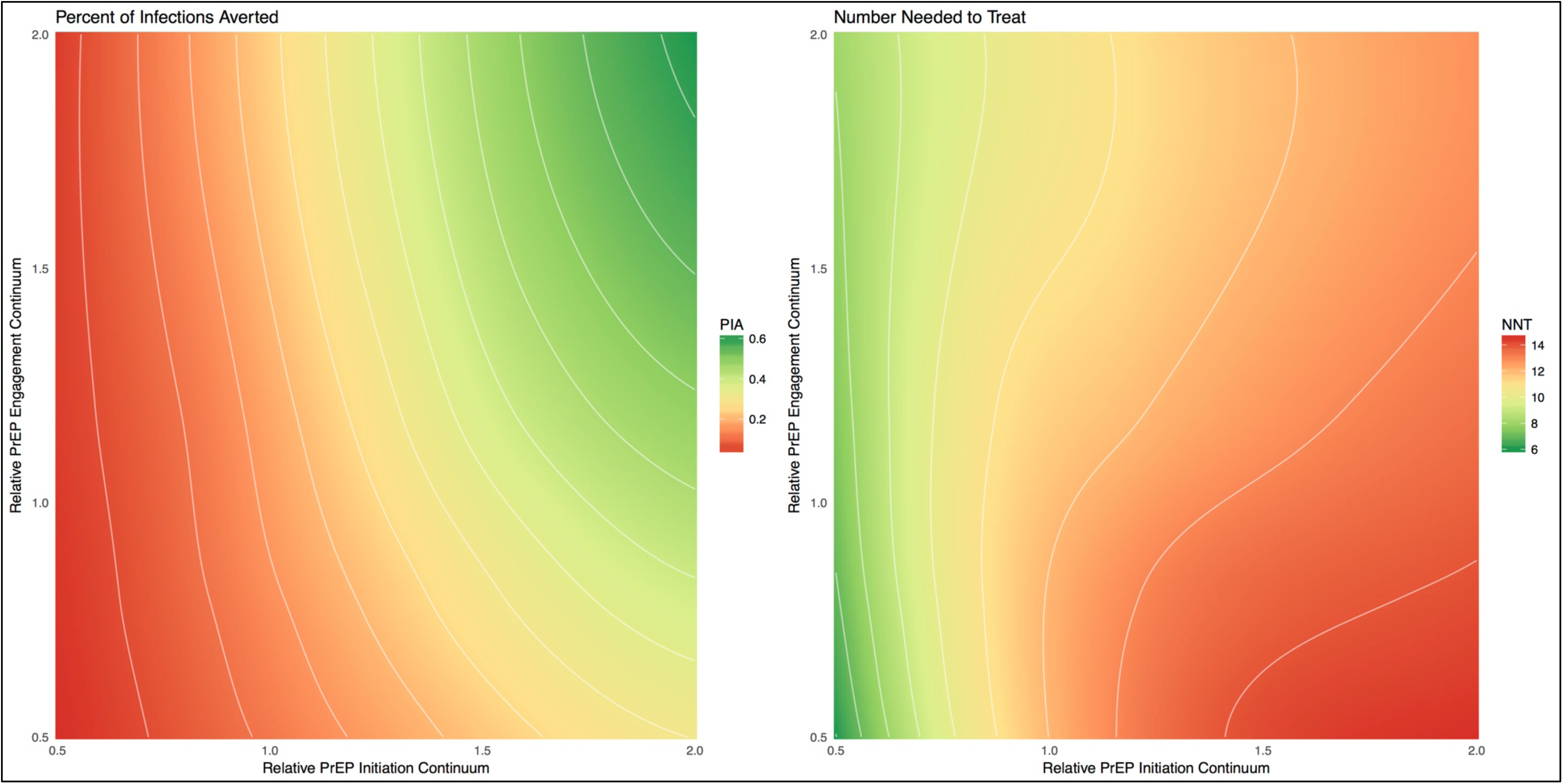
The percent of infections averted (PIA) for BMSM and number needed to treat on PrEP for one year to prevent one new HIV infection among BMSM across relative values of the combined BMSM PrEP continuum for initiation (factors = awareness, access, and prescription) versus engagement (factors = adherence and retention).

## DISCUSSION

In this modeling study, we found that implementation of PrEP could reduce absolute disparities in HIV incidence between BMSM and WMSM even despite current racial gaps in HIV PrEP care. Further disparity reduction with PrEP could be achieved with interventions targeting each of the modeled PrEP continuum steps for BMSM. Major gains in overall HIV incidence reduction and disparity elimination with PrEP would require targeting initiation factors (awareness, access, and prescription) over engagement factors (adherence and retention) for BMSM, given the currently observed PrEP continuum.

Many HIV prevention interventions successful at reducing HIV incidence are challenged by simultaneously addressing persistent HIV racial disparities. Systemic racial gaps in clinical care for testing and treatment of HIV (44) have led to a HIV prevention landscape in which white and higher-income MSM disproportionately benefit (45). We rooted our model structure and parameters in robust data to estimate how empirical representations of the PrEP continuum could impact HIV incidence over the next decade in a high-burden, low-resource population of younger BMSM (26). Our model suggests that it is possible to reduce, although not entirely eliminate, disparities in HIV incidence by race while at the same time lowering HIV incidence overall with PrEP.

To guide public health policy, we used a five-step PrEP continuum framework to conceptualize gaps in PrEP care (3). First, we found that awareness of PrEP was the step most strongly associated with incidence reduction for BMSM, partially due to the marginally declining conditional probabilities for the subsequent steps. Several studies have found reduced interest of BMSM in PrEP (12,13), related to lack of knowledge about PrEP and perceived stigma in using it (14,46,47). New technologies, such as mobile phone applications, are currently being developed to address this step. Second, PrEP access given awareness could increase infections averted by 4.1% if raised to observed WMSM levels in our model. Access-related interventions include patient assistance programs to cover medication costs (48); however, PrEP requires ongoing monitoring services covered through health insurance, which may be a barrier for some BMSM (15–17). Third, we found a relatively minimal effect for prescription rates conditional on access because the observed gap was only 10% (73% versus 63%). While all indicated BMSM seeking a PrEP prescription should receive one, this will depend on indications for PrEP being accurately queried by clinicians who are willing to prescribe PrEP. BMSM are less likely to be “out” to their doctors (49), and some clinicians may be less willing to prescribe to BMSM than to WMSM (50). Clinical training on PrEP patient assessment is greatly needed. Fourth, adherence is critical to both the impact and efficiency of PrEP, with a substantial effect on the NNT. Race/ethnicity has been strongly associated with suboptimal PrEP dosing (11,51). Long-acting formulations like injectable cabotegravir may benefit BMSM with adherence barriers (52). Finally, greater retention in PrEP care was strongly associated with both infections averted and lower NNT in our model. PrEP discontinuation for reasons other than lapsed indications has been an increasing challenge in clinical practice as PrEP users mature (53); lessons learned from managing patients with suboptimal levels of retention in HIV medical care may guide considerations of how to limit PrEP discontinuation (3).

We quantified disparities an absolute index that subtracts the standardized incidence rate of WMSM from the BMSM incidence rate, and a relative index that takes their ratio. Many policy documents use the latter: the National HIV/AIDS Strategy, for example, sets a goal to reduce racial disparities in new HIV diagnoses by a relative measure (6). In a dynamic intervention context, however, we would suggest that ratios are less suitable than differences for three reasons. First, the population-level burden of disease is quantified by the incidence rate of disease per unit of person-time. Using the absolute disparity allows one to express disparities with this same denominator. Second, there are parallels in using the absolute index with the choice of risk differences versus relative risks to quantify public health impact of a risk factor in epidemiological studies (54). Third, the ratio scale is unstable when the denominator is small relative to the numerator, as it is here. Ratio scales may be misleading for some interventional scenarios in these cases when reducing the difference in the number of incident infections between races has the counterintuitive effect of increasing the disparity ratio. Therefore, we recommend that disparities be quantified as absolute differences.

### Limitations

Our model conceptualizes racial disparities by simulating a two-race population of MSM of younger non-Hispanic black and white MSM in the Atlanta area. The conclusions drawn from this study are therefore most applicable to this target population. Deviations from random sampling of MSM in this target population from the two network/behavioral studies (11,21) could have resulted in biases in the estimates of model parameters in these domains. Specifically, because most parameters represent marginal probability and rate estimates, the resulting statistics in our models depend on the specific distribution of covariates in our particular study sample. An overrepresentation of young MSM in these studies, for example, could have resulted in upwardly biased behavioral risk parameters if positively correlated with age. Clinical and biological parameters were drawn from the secondary literature; aggregating multiple data streams into a single model requires strong assumptions about exchangeability, the implications of which have recently been examined in the methodological literature (55). However, a related strength of our study with respect to parameterization is its rigorous Bayesian model calibration and validation methods to evaluate and adjust for sources of parameter uncertainty through fitting the model projects to external HIV and STI prevalence and incidence data. Additionally, our model may be limited by the assumption that routine HIV screening is the primary point for entry into PrEP, based on the requirement that HIV testing be performed before PrEP initiation (37). Initiation of PrEP before specific sexual risk events has also been observed (10), and our future work will explore variations in reasons for starting PrEP. Finally, the continuum parameters were also based on two studies with race-stratified estimates, and these BMSM study populations may not represent other populations of HIV-uninfected BMSM in the US. Further parameter data are needed for other geographic settings to transport these findings to other MSM populations (56).

### Conclusions

PrEP will play a critical role in HIV elimination in the US, but its success will also depend on how we use it to address the persistent racial disparities in HIV incidence. Implementation of PrEP following a continuum framework could achieve the dual goals of reducing HIV incidence overall and decreasing the disparities in incidence between BMSM and WMSM over the next decade even despite current racial gaps in PrEP care. However, targeting these gaps with existing and novel interventions is greatly needed to make critical advances in using PrEP to reduce disparities.

## Acknowledgements

Author Affiliations. Emory University: Samuel M. Jenness, Kevin M. Maloney, Kevin M. Weiss, Patrick S. Sullivan; Centers for Disease Control and Prevention: Dawn K. Smith, Karen W. Hoover; University of Washington: Steven M. Goodreau, Darcy W. Rao; University of Albany: Eli S. Rosenberg; San Francisco Department of Public Health: Albert Y. Liu.

This work was supported by Centers for Disease Control [grant: U38 PS004646], the National Institutes of Health [R21 MH112449; R21 HD075662], and the Center for AIDS Research at Emory [grant: P30 AI050409] and the University of Washington [grant: P30 AI027757]. The authors declare no conflicts of interest.

We thank members of the scientific and public health advisory groups of the Coalition for Applied Modeling for Prevention project for their input on this study, and specifically those members who reviewed a previous version of this manuscript: Drs. Gregory Felzien and Jane Kelly.

The findings and conclusions in this paper are those of the authors and do not necessarily represent the views of the US National Institutes of Health or Centers for Disease Control and Prevention.

The authors have no conflicts of interest to declare.

